# Agronomic behavior of four cultivars and 56 F1 tomato populations

**DOI:** 10.1101/2021.04.03.438338

**Authors:** Gonzalo Quispe Choque

## Abstract

This research was conducted under a greenhouse at the National Vegetable Seed Production Center located in the municipality of Sipe Sipe, Quillacollo province with the aim of analyzing agronomic behavior in 56 F1 progenies and four hybrid tomato cultivars. The research consisted of 60 genotypes as treatments. Each experimental unit was made up of 6 plants. The variables evaluated were: number of flowers per inflorescence, number of fruits per cluster, weight of fruit, equatorial diameter, polar diameter, heart diameter, thickness of pericarp, number of fruits per plant, weight of fruit per plant, hardness and solids soluble . The data obtained were analyzed with descriptive statistics, correlation, cluster and principal component analysis. Significant correlations were found for the number of fruits per plant and number of flowers per inflorescence with the weight of fruit per plant. The polar dendogram identified four distant groups in the performance and quality components. The principal component analysis identified the degree of association between variables and genotypes, identifying groups correlated to components of fruit yield and quality.

## Introduction

Tomato (*Solanum lycopersicum* L.) is a diploid, self-pollinated species of the Solanaceae family with twelve pairs of chromosomes (2n = 24) considered an important crop that has a high nutrient content, is widely cultivated throughout the world and is the second most consumed vegetable after the potato (Mustafa, et al., 2017 and Chaudhari et al., 2019). In Bolivia, around 5,752 ha of tomato are cultivated, with a yield of 13 t ha-1. The highest percentages are provided by the mesothermic valleys of Santa Cruz (Saipina, San Isidro, Valle Grande and Los Negros) and valleys of Cochabamba (Omereque and Valle Alto) (MDRyT, 2013). Despite this, yields are low due to various factors, including pests and diseases, agronomic management, and market prices. These factors constitute the main limitations for tomato production in Bolivia, significantly reducing production levels, with an increasingly frequent risk of complete losses (Crespo, 2007).

One way to mitigate this problem is to obtain varieties resistant to pests and diseases with high yield potential, through genetic improvement. Unfortunately, the cultivated tomato has experienced severe genetic bottlenecks through the domestication and selection process, in addition to pollination by autogamy (Rick, 1976; Bai and Lindhout, 2007 and Torrico et al., 2015), these two factors have combined so that the tomato genome is very narrow genetically and has very little genetic diversity (Miller and Tanksley, 1990).

The foregoing has led genetic improvement programs to form base populations from crosses between lines, from crossing between improved varieties, by crossing a group of lines or populations (gene pool) or a native population per improved variety (Dzib-Aguilar et al., 2011). Given the context, the INIAF Vegetable Project has directed activities to the formation of segregating populations with the use of germplasmic resources, seeking to respond to the almost null existence of national varieties in the country, developing this work with the objective of analyzing the agronomic behavior in 56 F1 progeny and four hybrid tomato cultivars.

## Materials and methods

The material used in this research was 56 direct crosses of 20 parental lines selected from the working collection, which were evaluated for their attributes of performance components and disease resistance. The crosses were made based on the factorial genetic design or design II, which consists of crossing between m male parents and with a group of h female parents to obtain mh families. The formation of the hybrids was carried out in a greenhouse of the National Center for Vegetable Seed Production (CNPSH), Villa Montenegro locality, Quillacollo province, geographically located at 17 ° 22 ‘S; 66 ° 19′W, at an altitude of 2505 masl, during the 2018-2019 agricultural season. This area has an average temperature of 23 ° C. The fruits from each cross were harvested at the physiological maturity stage and stored until their complete maturity, in order to extract the seed. The evaluation of the 54 progenies was carried out in the 2018-2019 agricultural season at the same Innovation Center.

The sowing of the progeny and hybrid cultivars (Lia, Santy, Shaman and Bingo) was carried out in 128-cell polyethylene trays, containing prepared substrate at a 1: 1: 1 ratio of rice husk, topsoil and fine sand, which It was disinfected in a steam boiler for 45 min at 90 ° C, sowing 15 seeds of each genotype, applying light irrigation and later they were placed in the greenhouse for their germination and development. The transplant was carried out in a greenhouse in black 400 micron polyethylene sleeves under a staggered system. The experimental unit consisted of six plants at a distance of 40 cm between plants. After the transplant, the irrigation application was started at a rate of three times a day for a period of 20 min, increasing the irrigation time according to the needs of the plant.

Variables associated with fruit yield components of the second cluster were evaluated in three individual plants per plot, in complete competition. To estimate the average weight of the fruits in each material, the weight of each of the harvests carried out was added and divided by the number of total fruits. At the third harvest, 5 random fruits of each genotype were taken, their individual weight, polar diameter, equatorial diameter of the fruit, pericarp thickness, heart diameter, number of locules and fruit shape were taken, at the end the average was reported. in the quantitative variables and mode in the qualitative variables of the 5 fruits; These were evaluated according to the descriptor manual for tomato (S. lycopersicum L.) of the International Plant Genetic Resources Institute (IPGRI,

Performance and its components were analyzed using descriptive statistics andPhenotypic correlations (r) between the variables, using the Pearson correlation coefficient, a dendrogram was generated using the 13 quantitative and qualitative variables, the genetic distances were calculated using the Average method and the Eucledian distance, the variation between progeny was also analyzed. by main components.The data processing and statistical analysis was carried out with the program R Project 3.2.5 (2016).

## Results and Discussion

The phenotypic data of the genotypes are shown in Table 1 based on the results of descriptivite statistics. With these statistics, useful information is obtained that allows inferring several results and questions, for example, the number of fruits per plant and their weight have a CV> 50%, whichpoints outwhich has the highest variability. Likewise, nine variables appear with CV> 10%, which indicates that genotypes may have little variability in these characters. In the same way, the dispersed behavior is observed in the fruit weight with an average of 84.6 g.González and Laguna, (2004) cited by Blandon (2017) affirm that the differences in fruit weight between the genotypes are due to the genetic makeup of each line and the influence exerted by the environment.

**Table 1.** Descriptive statistics for eleven quantitative characters evaluated in nine experimental tomato lines during the agricultural season 2017-2018.

| Variable | Half | Des. Its<br>T. | Minimum | Maximum | Rank | CV<br>% |
| --- | --- | --- | --- | --- | --- | --- |
| **Performance components** |  |  |  |  |  |  |
| Number of flowers per inflorescence (Units) | 5.08 | 0.66 | 4 | 6.5 | 2.5 | 13 |
| Number of fruits per bunch (Units) | 3.61 | 1.06 | 3 | 6.5 | 5.5 | 27.7 |
| Fruit weight (g) | 84.6 | 22.7 | 47.1 | 155 | 108.30 | 26.8 |
| Equatorial diameter (mm) | 48.9 | 5.1 | 37.7 | 63.5 | 25.81 | 10.4 |
| Pole diameter (mm) | 56.1 | 6.47 | 40 | 73.2 | 33.20 | 11.5 |
| Heart diameter (mm) | 23.3 | 4.09 | 15.8 | 37.6 | 21.80 | 17.6 |
| Pericarp thickness (mm) | 7.4 | 0.99 | 5.25 | 9.92 | 4.67 | 13.4 |
| Number of fruits per plant (Units) | 21.1 | 12 | 6 | 82.5 | 76.5 | 57 |
| Fruit weight per plant (g) | 1447 | 725 | 381 | 3643 | 3261.65 | 50.1 |
| **Quality Parameters** |  |  |  |  |  |  |
| Hardness (Shore) | 69.4 | 9.78 | 43.8 | 93.6 | 49.80 | 14.1 |
| Total soluble solids (° Brix) | 7.07 | 0.8 | 5.75 | 10.9 | 5.10 | 11.3 |

The results of the phenotype correlation showed that a total of 26 correlations were significant (p <0.05) (Figure 1). Equatorial diameter, polar diameter, heart size and pericarp thickness were found to have a positive correlation with fruit weight. While the pericarp thickness was correlated with the weight of the fruit, polar and equatorial diameter. Another important association was formed by the number of flowers per inflorescence with the number of fruits per cluster (r = 0.53) and the weight of the fruit per plant (r = 0.41). Bustamante (2014) concludes that the quantity of fruits are defined by the genetic characteristics of varieties and hybrids of this crop. Similarly, Zárate (2007) when evaluating the number of fruits per plant of hybrid tomatoes in the hydroponic system and soil, obtained different amounts of fruits per plant. Another important correlation is given by the pericarp thickness with the weight, polar and equatorial diameter of the fruit (r = 0.55, r = 0.38, r = 0.48). A high correlation was also observed between the weight of fruit per plant and the number of fruits per plant.

**Figure 1.**
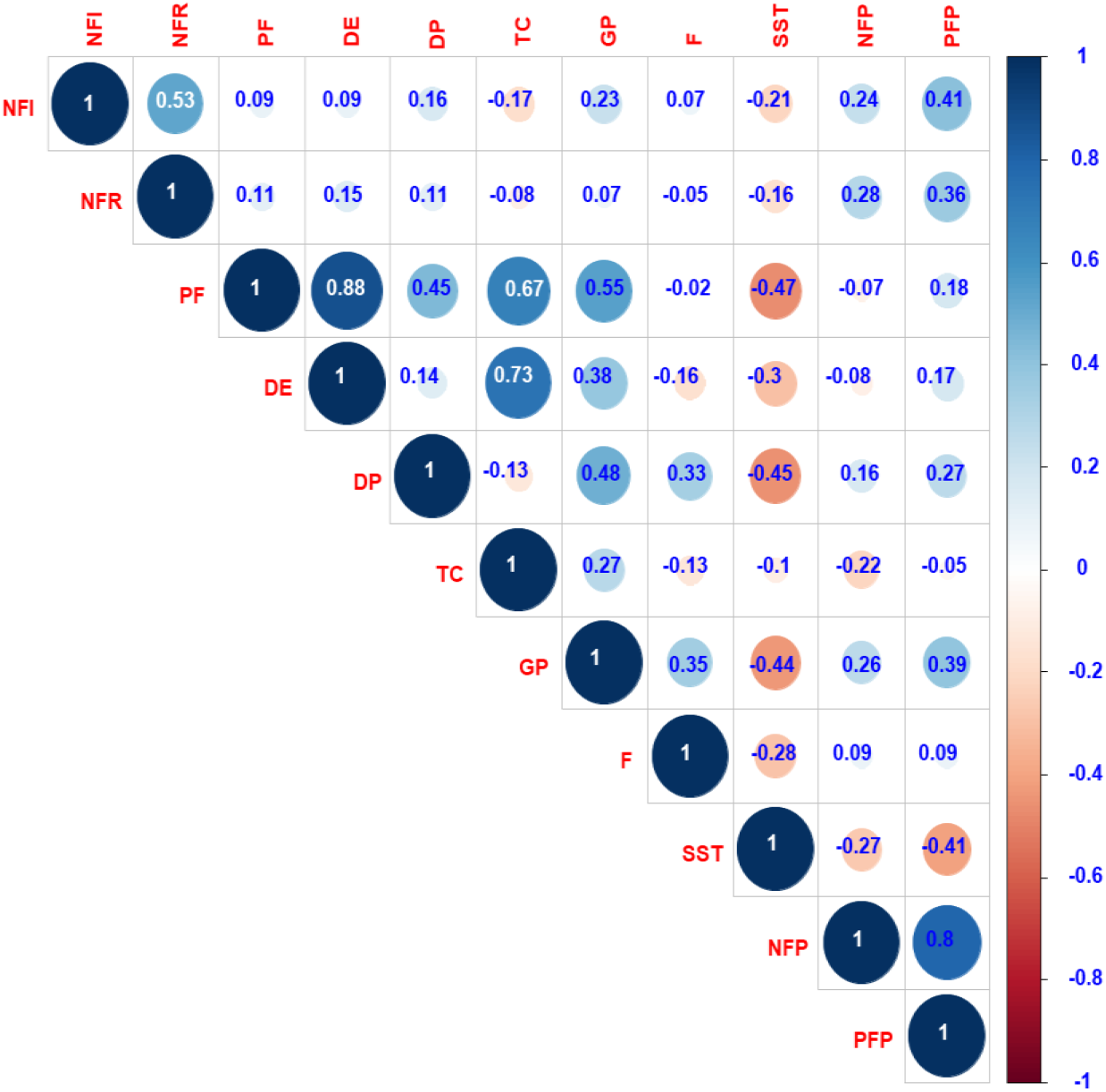
Correlogram of the degree of association between the yield of the fruit and its components in experimental tomato lines, evaluated during the 2018-2019 agricultural season in Villa Montenegro, Sipe Sipe, Cochabamba.

The polar dendrogram of Figure 2 obtained through the Eucledian distance shows the formation of four groups at a cutoff of 0.6, these were grouped according to characteristics of similarity between genotypes. In group I, the genotypes with the greatest relationship between equatorial and polar diameter were located. In addition, they had the lowest number of fruits per bunch, unlike the other groups. Group II included progeny with pyriform fruits as they had greater polar than equatorial diameter and developed fewer fruits per plant and pericarp thickness. The fruits of these materials resemble the hybrid cultivars Xaman, Santy and Bingo included in this group, presenting high values in quality characteristics, a character that is important for researchers. In group III they found a greater number of flowers per inflorescence, with high values in polar diameter, pericarp thickness, fruit weight and hardness. According to Ramos et al. (2010), this last characteristic is considered important for the fresh tomato commercialization. In group IV, progeny 125 and 132 were located, whose inflorescences had a greater number of fruits per inflorescences and fruits per plants.

**Table 2.** Comparison of Duncan means for nine quantitative characters evaluated in nine experimental tomato lines during the 2017-2018 agricultural season.

| Variable | Conglomerate |  |  |  |
| --- | --- | --- | --- | --- |
|  | G1 | G2 | G3 | G4 |
| **Performance components** |  |  |  |  |
| Number of flowers per inflorescence (Units) | 5.18 | 5.04 | 6 | 5 |
| Number of fruits per bunch (Units) | 3.71 | 3.68 | 4.46 | 5 |
| Number of fruits per plant (Units) | 23.24 | 13 | 33.92 | 40.5 |
| Fruit weight per plant (g) | 1564.93 | 831.33 | 2322.69 | 3384.28 |
| Fruit weight (g) | 86.91 | 80.43 | 89.83 | 88 |
| Equatorial diameter (mm) | 50.3 | 47.55 | 49.72 | 49.01 |
| Pole diameter (mm) | 53.39 | 55.49 | 60.47 | 58.22 |
| Heart diameter (mm) | 25.01 | 22.82 | 21.71 | 25.03 |
| Pericarp thickness (mm) | 7.5 | 7 | 8.09 | 7.65 |
| **Quality parameters** |  |  |  |  |
| Hardness (Shore) | 69.39 | 68.37 | 71.42 | 71.35 |
| Total soluble solids (° Brix) | 7.08 | 7.34 | 6.51 | 6.98 |

**Figure 2.**
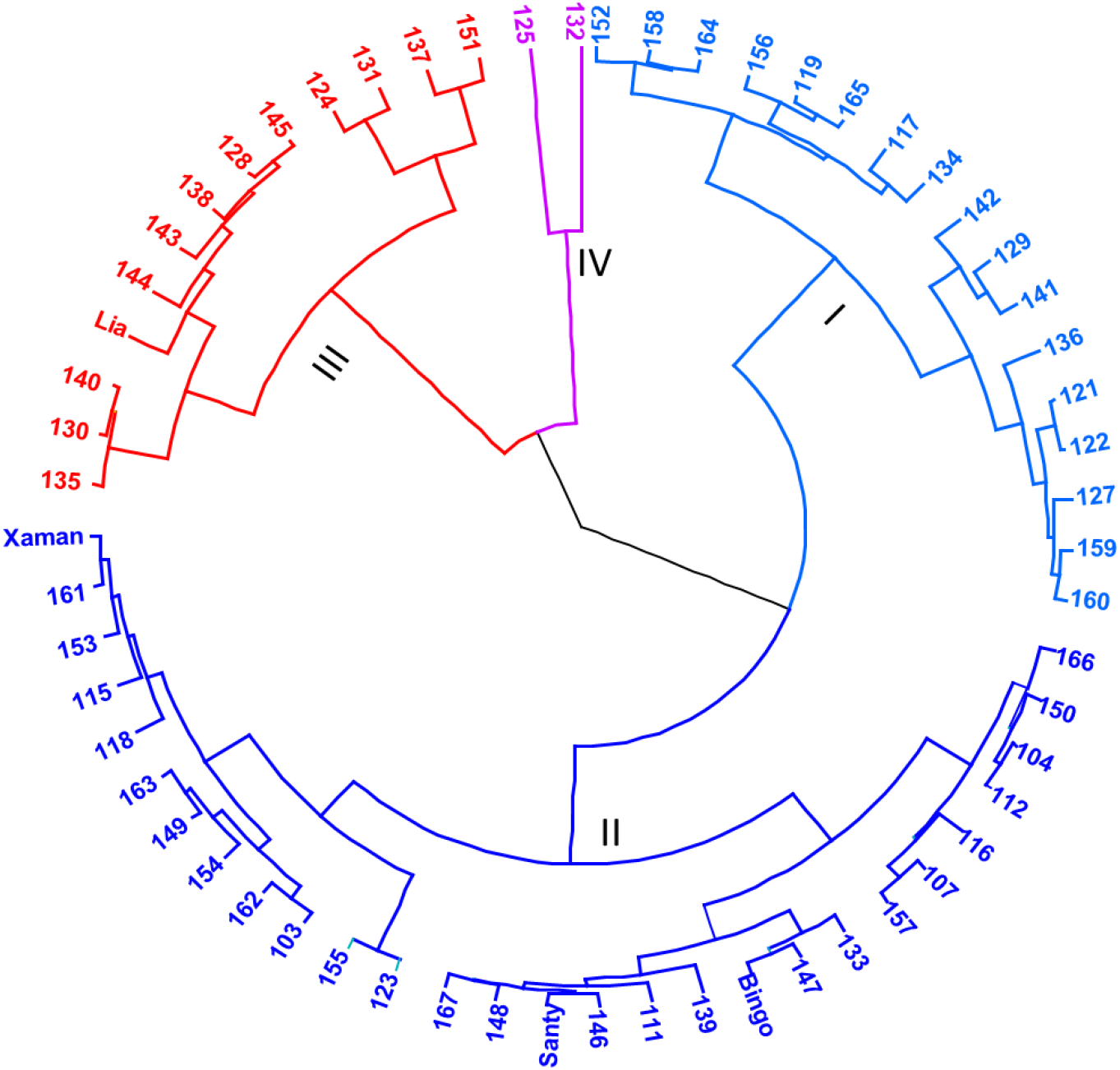
Polar dendrogram with variables of yield and quality components in 60 tomato genotypes, evaluated during the 2018-2019 agricultural season in Villa Montenegro, Sipe Sipe, Cochabamba.

With the purpose of transforming correlated variables into uncorrelated principal components, with independent interpretation one of the other. The contribution of the variables in the first two components is expressed with a value of 54.82% of the accumulated variance. The graphical representation of the main components of Figure 3, synthesizes the behavior of the genotypes with respect to the variables under study. The interpretation was made considering the magnitude of the vector, the direction and the angle they form with each other. Where it can be seen that the variables most positively linked to the first axis are pericarp thickness (GP), fruit length (LF), fruit weight per plant (PFP) and in a negative way total soluble solids. The variables most linked to the second axis are in a positive sense the number of fruits per plant (NFP) and in a negative sense the size of the heart (TC). The opposite projection of total soluble solids on the first axis in relation to the yield component variables means that the genotypes develop less solid content as fruits with greater pericarp thickness and polar diameter are obtained. The degree of association between variables and genotypes can also be observed, identifying groups correlated to components of fruit yield and quality. In this described association, the location of genotypes 130, 125, 137,131, 132, 143, 124, 140, 144, 138, 151 and Lia stands out. The opposite projection of total soluble solids on the first axis in relation to the yield component variables means that the genotypes develop less solid content as fruits with greater pericarp thickness and polar diameter are obtained. The degree of association between variables and genotypes can also be observed, identifying groups correlated to components of fruit yield and quality. In this described association, the location of genotypes 130, 125, 137,131, 132, 143, 124, 140, 144, 138, 151 and Lia stands out. The opposite projection of total soluble solids on the first axis in relation to the yield component variables means that the genotypes develop less solid content as fruits with greater pericarp thickness and polar diameter are obtained. The degree of association between variables and genotypes can also be observed, identifying groups correlated to components of fruit yield and quality. In this described association, the location of genotypes 130, 125, 137,131, 132, 143, 124, 140, 144, 138, 151 and Lia stands out. identifying groups correlated to components of fruit yield and quality. In this described association, the location of genotypes 130, 125, 137,131, 132, 143, 124, 140, 144, 138, 151 and Lia stands out. identifying groups correlated to components of fruit yield and quality. In this described association, the location of genotypes 130, 125, 137,131, 132, 143, 124, 140, 144, 138, 151 and Lia stands out.

**Figure 2.**
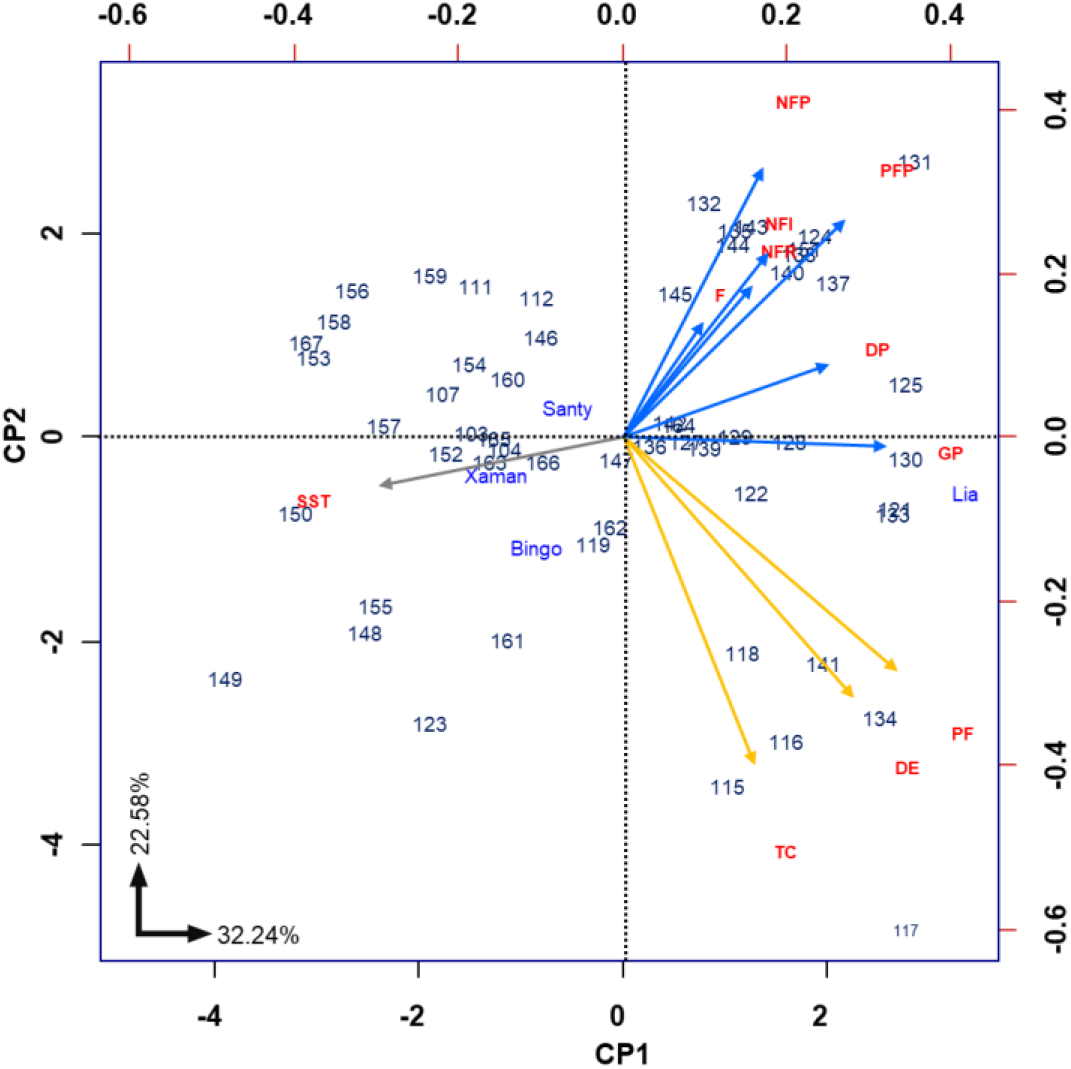
Principal component analysis for yield and quality component variables of 60 tomato genotypes, evaluated during the 2018-2019 agricultural season in Villa Montenegro, Sipe Sipe, Cochabamba.

## Conclusions

The variables of yield and quality components of the fruit were the most important characteristics that allowed the formation of groups, in which it is suggested to emphasize to lay foundations in the selection criteria, which are based on the correlation between they.

Due to their agronomic characteristics, most of the genotypes of group III and IV have potential in the productivity and quality components, which are similar to the characteristics of the control (Lia) and some of them could be used as a source of germplasm for development. of triple and double crosses in the genetic improvement program of the Vegetable Project.

## Cited References

UPOV (2011). International Union for the Protection of New Varieties of Plants. Tomato (Solanum lycopersicum L.). Guidelines for the execution of the examination of distinction, uniformity and stability. Document TG / 44/11.

IPGRI. (nineteen ninety six). International Institute of Plant Genetic Resources. Descriptors for Tomato (Lycopersicon spp.). International Institute of Plant Genetic Resources. Rome Italy. 49 p.

Marlina Mustafa, Muhamad Syukur, Surjono Hadi Sutjahjo and Sobir. (2017). Inheritance of Fruit Cracking Resistance in Tomato (Solanum lycopersicum L.). Asian Journal of Agriculturalb Research. 11(1), 10–17. 10.3923/aiar.2017.10.17

Ramos GM, SB Bautista, NL Barrera, ME Bosques and CM Estrada (2010). Antimicrobial compounds added to edible coatings for use in fruit and vegetable products. Mexican Journal of Phytopathology 28, 44–57.

Bai Y. and P. Lindhout (2007) Domestication and breeding of tomatoes: What have we gained and what can we gain in the future?. Annalsof Botany 100, 1085–1094

Rick CM (1976) Toma to, Lycopersicon esculentum (Solanaceae). In: Evolution of Crop Plants, NW Simmonds (ed). Ed. Longman Group. London. pp: 268–273.

Miller JC and SD Tanksley (1990). RFLP analysis of phylogenetic relationships and genetic variation in the genus Lycopersicon. Theoretical and Applied Genetics 80, 437–448.

Torrico, A., Crespo, M., Rojas, J., (2015). Morphological and molecular study of the genetic diversity of Bolivian wild tomato (Solanum spp.). Agriculture Magazine. 55, 20–28.

Crespo M. (2007) Analysis of tomato production in Bolivia. In a research proposal on “Development of differentiated products with added value in tomatoes and peppers, appropriate for a sustainable family production ex the Southern Cone” FONTAGRO.

Dzib-Aguilar, LA; Segura, JC; Ortega, R. and Latournerie, L. (2011). Diallelic crosses between native yucatan maize populations and improved populations. Trop. Subtrop. Agroecosys. 14: 119–127.

Bustamante, N. (2004). Adaptability of four varieties of kidney tomato lycopersicum sculentum mill, Cango site, Putuyango Canton. Obtained fromhttps://dspace.unl.edu.ec:https://dspace.unl.edu.ec/jspui/bitstream/123456789/5573/1/Rengel=20Bustamante=20Nelson.pdf

Zárate, B. (2007). “Production of hydroponic tomato (lycopersicon esculentum Mill.) With substrates, under greenhouse”. Obtained fromhttp://tesis.ipn.mx:http://tesis.ipn.mx/jspui/bitstream/12345789/779/1/TESIS_MAESTRIA_BALDOMERO.pdf

